# A Curated Genome-Scale Nucleotide Diversity Panel of Non-Human Primates

**DOI:** 10.64898/2026.06.16.732573

**Authors:** Vasili Pankratov, Bjarke Meyer Pedersen, Erik Fogh Sørensen, Kasper Munch, Thomas Bataillon, Mikkel Heide Schierup, Juraj Bergman

**Author notes:** Authors contributed equally (Vasili Pankratov, Bjarke Meyer Pedersen).

## Abstract

**Background:** Primates constitute one of the most phylogenetically and ecologically diverse Eutherian mammalian orders, with a central role in advancing our knowledge of human evolution, speciation processes and conservation biology. While thousands of whole-genome sequences have been generated across a multitude of primate taxa, discrepancies in data processing - particularly the lack of ploidy-aware variant calling in sex-linked regions - have limited the utility of existing datasets for large-scale comparative analyses.

**Results:** Here, we utilized publicly available short-read sequencing data of non-human primates, recently published primate genome assemblies and a ploidy-aware variant calling procedure to generate a genome-scale nucleotide diversity panel comprising 3,240 individuals from 269 species and 71 genera. To further facilitate cross-species comparisons, we generated a multiple-genome alignment of primate assemblies used for variant calling.

**Conclusion:** This curated resource of non-human primate diversity provides a foundation for future research in primate evolutionary biology, speciation, and sex chromosome evolution (https://pure.au.dk/portal/en/datasets/primate-diversity-panel/).

## Background

Primates are one of the most speciose orders of mammals, comprising over 500 distinct species that span a wide range of ecological niches and geographic regions, primarily across tropical and subtropical zones in Africa, Asia, and the Americas^1^. Given that humans are part of the primate lineage - and widespread interest in our own evolutionary origins - the genomic sequencing of our closest primate relatives is essential not only for understanding human evolution but also for advancing the broader fields of comparative genomics and evolutionary biology. To date, whole-genome datasets have been generated for thousands of non-human primate individuals spanning the majority of primate genera^2–4^.

Despite the growing availability of genomic resources, comparative studies across primates remain challenging due to inconsistencies in data processing and a general lack of coordinated efforts to standardize nucleotide variant calling across datasets. In particular, ploidy-aware variant calling - a built-in feature of most state-of-the-art variant-calling programs^5^ - has been largely overlooked in the processing of primate genomic data. These limitations continue to restrict the broader utility of existing datasets for comparative genomic analyses across the primate lineage.

Here, we present a genome-scale nucleotide diversity panel of non-human primates comprising 3,240 individuals that cover 269 species and 71 genera, generated using publicly available sequencing datasets. Where possible, we prioritized sequencing data from populations that reflect natural genetic variation, focusing on individuals sampled from wild populations while excluding those sequenced as part of biomedical or clinical research studies. Additionally, our bioinformatic workflow utilized the latest primate genome assemblies (including seven telomere-to-telomere assemblies)^6,7^ for variant calling and takes into account the ploidy of sex-linked regions with respect to the sex of the individual. This resource (https://pure.au.dk/portal/en/datasets/primate-diversity-panel/) thus provides a valuable foundation for future studies in primate evolution and conservation, sex chromosome biology and comparative genomics.

## Results and Discussion

### Sample statistics

We constructed a comprehensive genomic resource comprising 3,240 samples, with the aim of creating the most extensive comparative datasets for non-human primates to date (Supplementary Table S1). This collection spans 269 species across 71 genera sourced from 148 unique BioProjects. The dataset includes 2,056 Old World monkeys (*Cercopithecidae*), 526 apes (*Hominoidea*), 331 New World monkeys (*Platyrrhini*), 322 strepsirrhines (*Strepsirrhini*), and five tarsiers (*Tarsiidae*). Through genetic sex determination^8^ (see Methods; Supplementary Fig. S1), we substantially improved sample annotation, reducing the number of samples with unknown sex from 1,384 to five, resulting in a final dataset of 1,756 females (previously 1,036) and 1,479 males (previously 820). The five individuals of unknown sex, which are the five tarsier individuals, were treated as female in all downstream analyses. The average read coverage of samples in our dataset (defined as the average read depth within regions of the reference sequence covered by at least one mapped read) was 22.31× genome-wide, 22.57× for the autosomes (Fig. 1) and 17.53× for the X chromosome. The average fraction of the covered reference genomes (defined as the proportion of the reference sequence covered by at least one mapped read) was 83% genome-wide, 84% for the autosomal (Fig. 1) and 81% for the X-linked contigs, across samples. The proportions of samples with at least 10× average read coverage across the autosomal and X portions of the reference genomes were 63% (2,027 individuals; Fig. 1) and 59% (1,902 individuals), respectively.

**Figure 1.**
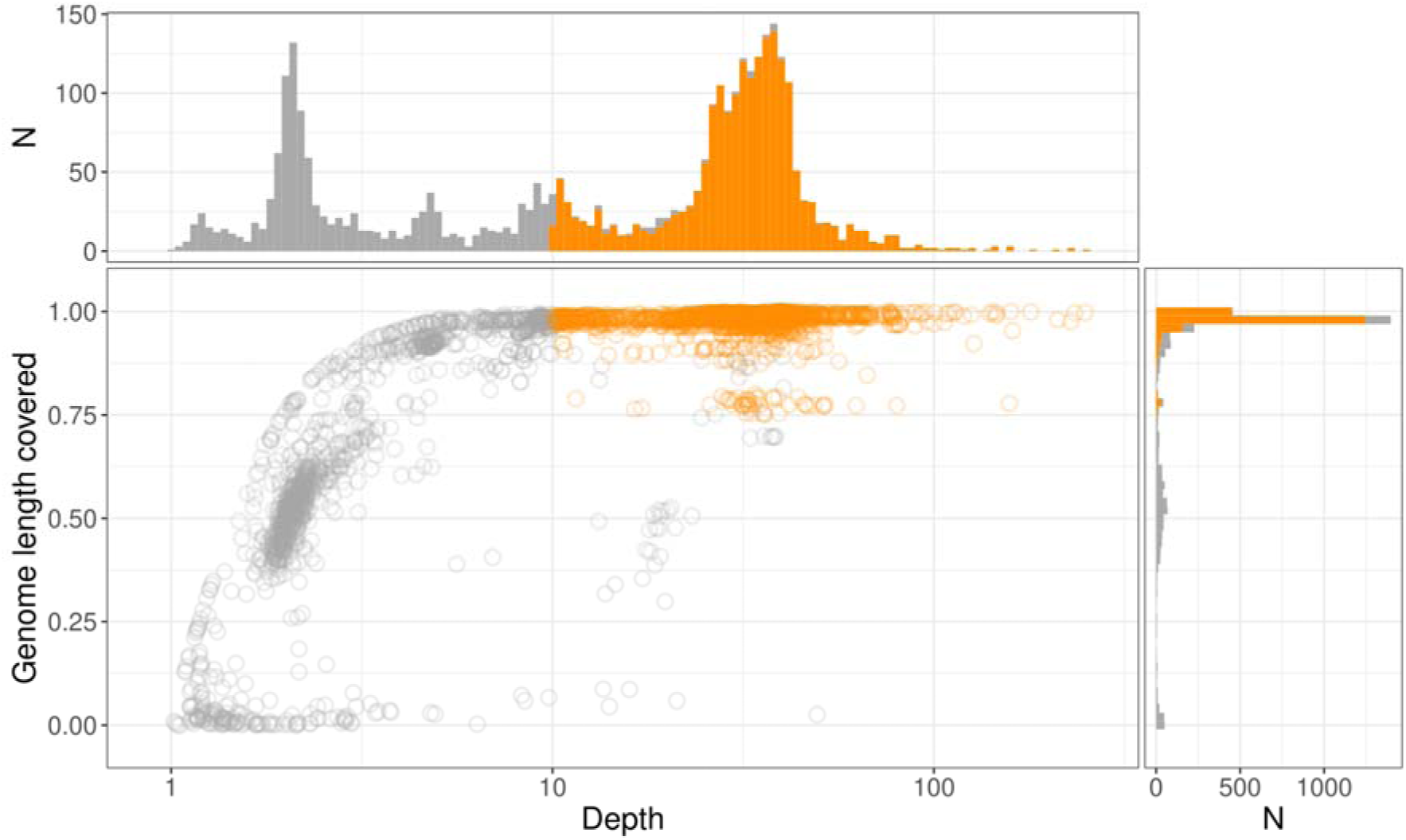
Autosomal sequencing depth (total length of mapped reads divided by the length of the autosomal portion of the reference genome with at least one read mapping to it) and fraction of autosomal portion of the reference genome with at least one read mapping to it for each sample. For each reference genome, only autosomal contigs equal to or longer than 1 Mb were considered. Samples shown in orange have sequencing depth equal to or greater than 10× and fraction of the genome covered equal to or greater than a species-specific cut-off threshold specified in Supplementary Table S1. These samples were used to define the species-specific callability masks. Panels above and on the right show sample distributions with respect to sequencing depth and fraction of the covered genome, respectively.

The genome-average read mapping quality was above 40 in 3,164 samples and above 50 in 2,728 samples, while the fraction of mapped reads is above 50% in 3,166 samples and above 75% in 3,100 samples (Supplementary Fig. S2A). Most samples with low read mapping quality or low fraction of mapping reads were also characterised by low coverage and generally low quality (see Sample Quality Control in Methods; Supplementary Fig. S2B). Indeed, out of 92 samples with mapping quality below 40 and/or fraction of mapped reads below 50%, 32 came from bioproject PRJNA508503, which used urine or faeces as DNA source; 19 from PRJNA16955, which did a *de novo* assembly and included sequences of specific clones used for scaffolding as separate biosamples; 17 from PRJEB77609, which likely had a sample labelling error (SRA data from this project is no longer available from NCBI); and 12 from PRJEB60463, which used a single hair as DNA source. On the other hand, while high sequence divergence from the reference genome resulted in a lower fraction of mapped reads, it had a modest effect on mapping quality.

### Reference genome alignment

To enable comparative analyses between species, we used the Progressive Cactus toolkit^9^ to produce a multiple-genome alignment of 49 reference assemblies, comprising the 47 non-human primate genomes used for short-read mapping, as well as the *Homo sapiens* reference assembly and the *Tupaia belangeri* outgroup (Supplementary Table S2; Supplementary Fig. S3). The alignment includes a total of eight telomere-to-telomere (T2T) gap-free references, including the human T2T-CHM13v2.0 reference^10^, the crab-eating macaque (*Macaca fascicularis*) T2T reference^6^, as well as five great ape and the siamang (*Symphalangus syndactylus*) T2T references^7^. For each non-human reference sequence, Fig. 2 shows the fraction of the total length of the human genome that is alignable to every other reference sequence, with this fraction decreasing with increasing phylogenetic distance. Conversely, the number of mismatches between the human genome and other references increased with phylogenetic distance, as expected. Notably, the divergence between human and non-human X chromosomes was lower compared to the divergence of autosomal sequences.

**Figure 2.**
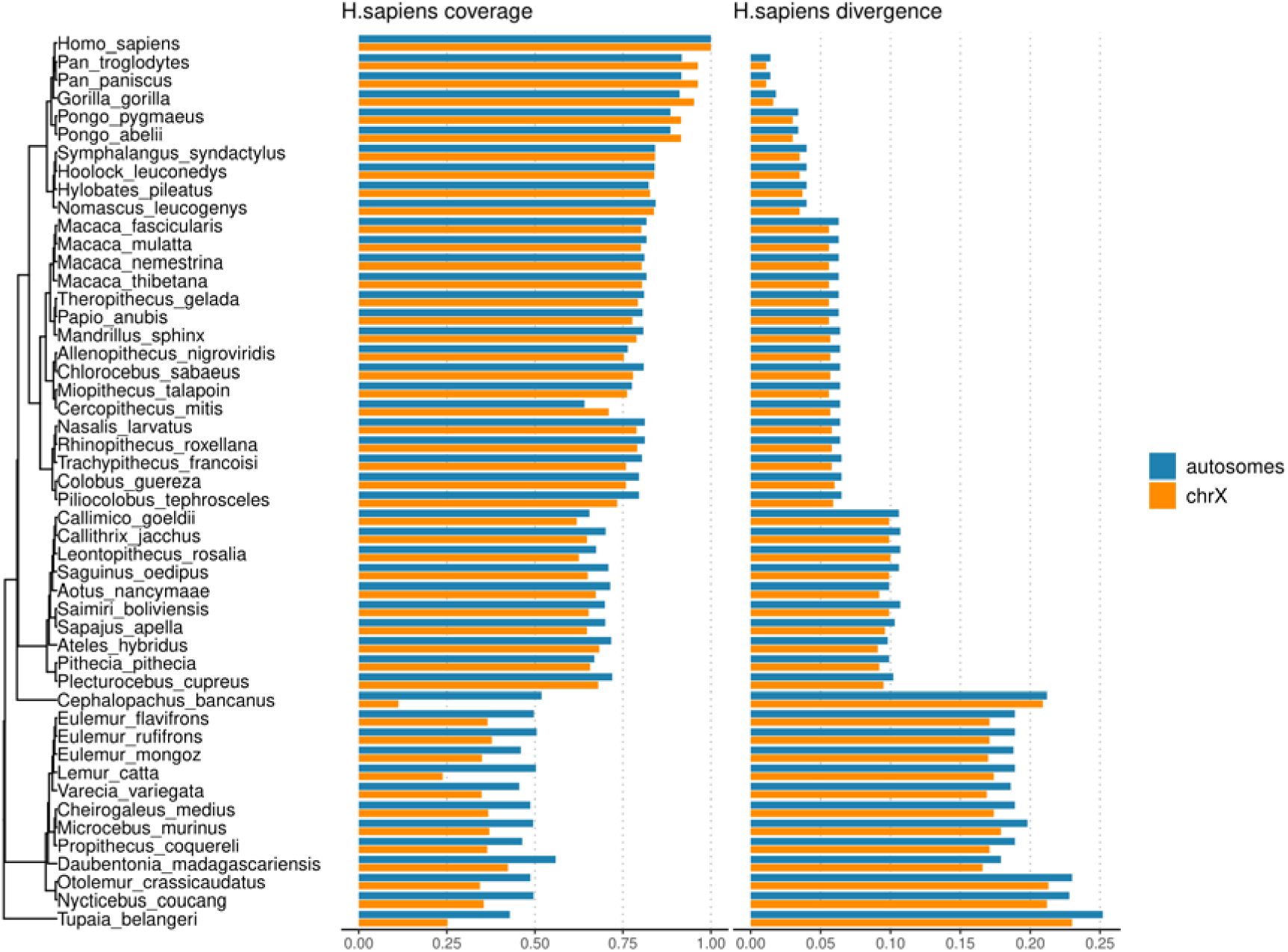
Comparison of reference genomes to the T2T-CHM13v2.0 human genome. Coverage refers to the fraction of the total length of the human genome with sequence from the query genome mapping to it (not necessarily 1-to-1). Divergence refers to the number of mismatches between the human and query sequence, normalised by the length of the aligned sequence. Both statistics are presented separately for human autosomes and human chromosome X.

We also harnessed the multiple sequence alignment to identify contigs derived from the X chromosome based on their homology to the human X chromosome. This allowed us to annotate X-chromosome-derived contigs in 23 reference genomes for which this information was not available, as well as correct an erroneous chromosome X annotation in the *Piliocolobus tephrosceles* genome (Supplementary Table S3; Supplementary Fig. S3).

### Nucleotide diversity analysis

The following analyses were restricted to individuals passing quality control including autosomal mapping depth equal to or greater than 20× and genotype missingness proportion < 1%, resulting in a total of 1,427 indivudals across 255 species and 71 genera (Fig. 3A). We calculated autosomal and X chromosome individual-based heterozygosity across primate families and observed a large variation in autosomal heterozygosity amongst groups (Fig. 3B), with median values ranging from 1.5×10□³ in New World monkeys to 2.8×10□³ in tarsiers, while the X chromosome varied from a median of 3.5×10□□ in apes to 8.15×10□□ in strepsirrhines (Fig. 3C). For the majority of samples, the X:A heterozygosity ratio was below 0.75 (Fig. 3D), indicating that the majority of primate species were affected by processes that disproportionately reduced X chromosome diversity compared to autosomal diversity. Such processes may involve male-biased migration^11^ and mutation processes^12–14^, as well as intensified selective pressures on X-linked variants across primates^15^, among other factors^16–19^.

**Figure 3.**
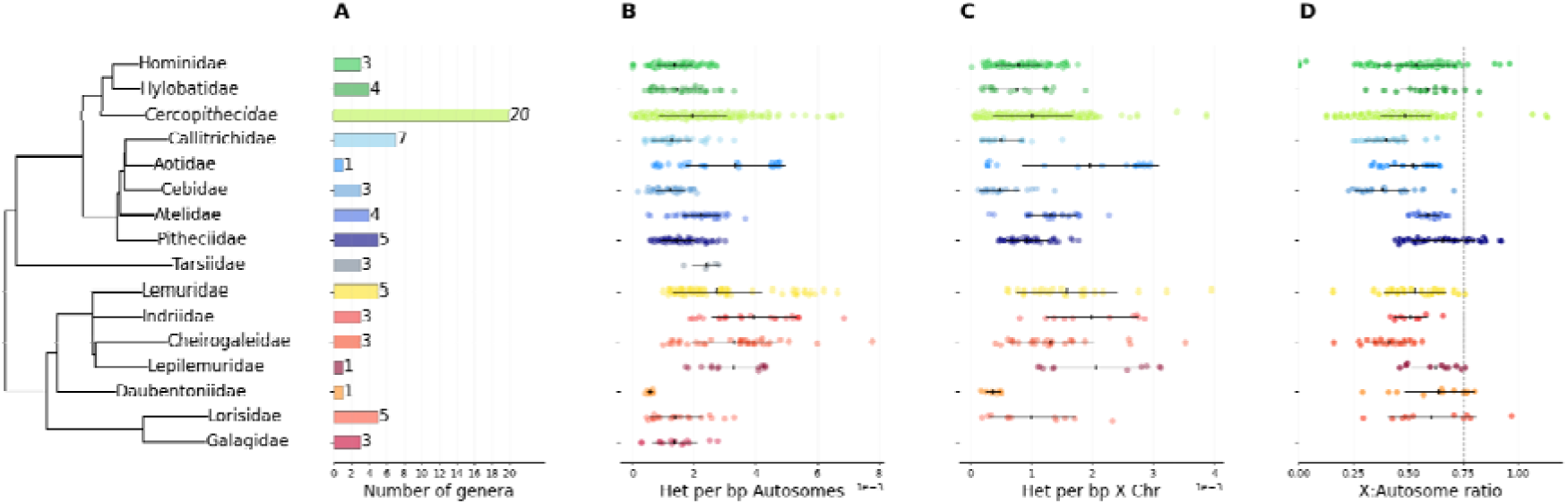
Patterns of genomic diversity across 16 primate families. **A.** Number of genera represented within each primate family included in the dataset. **B.** Autosomal heterozygosity per base pair across individuals. **C.** X chromosome heterozygosity per base pair across individuals. **D.** Ratio of X chromosome to autosomal heterozygosity. Horizontal black bars indicate mean values per family, with error bars representing standard error.

We next performed principal component analysis (PCA) on genetic variants for large subsamples of individuals within our dataset, comprising individuals mapped to the *Papio anubis* reference (Fig. 4 A, B; 251 individuals) and those mapped to the *Macaca mulatta* reference (Fig. 4 C, D; 183 individuals). Autosomal and X-chromosome variant-based PCAs yielded similar clustering patterns. In *Papio* species, PC1 separated northern taxa (*P. anubis*, *P. hamadryas*, and *P. papio*) from southern taxa (*P. cynocephalus*, *P. kindae*, and *P. ursinus*), while PC2 distinguished *P. hamadryas* and *P. kindae* from the larger *P. anubis* and *P. cynocephalus* clusters. Several individuals known to be hybrids between *P. anubis* and *P. cynocephalus* fell between the primary *P. cynocephalus* and *P. anubis* clusters. One individual labelled as *P. kindae* clustered with *P. anubis*, suggesting a probable species mislabeling. Additionally, *P. cynocephalus* individuals known to carry mixed *P. kindae*–*P. cynocephalus* ancestry clustered more closely with *P. kindae*, compared to the primary *P. cynocephalus* cluster.

**Figure 4.**
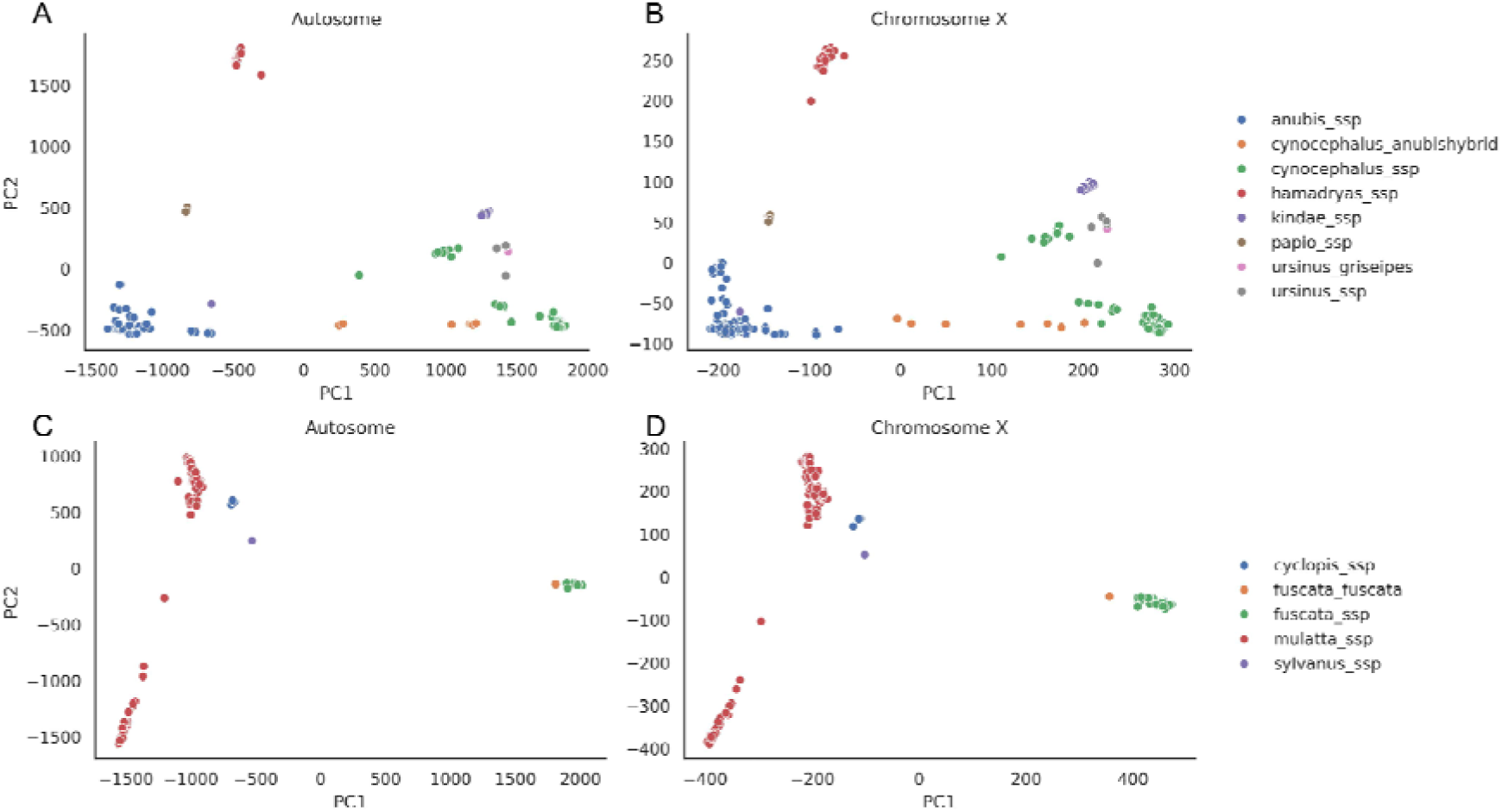
PCA of individuals mapped to *Papio anubis* and *Macaca mulatta* reference assemblies. **A.**, **C.** PCA based on the autosomal variants for *Papio* and *Macaca* species, respectively. **B.**, **D.**) PCA based on the X-linked variants for *Papio* and *Macaca* species, respectively.

The four macaque species aligned to the *M. mulatta* reference (*M. mulatta*, *M. fuscata*, *M. sylvanus*, and *M. cyclopis*) had vastly different sample sizes, with only a few samples from *M. sylvanus* and *M. cyclopis* species. PC1 split *M. fuscata* from the other species, while *M. mulatta* was split into two distinct clusters on PC2, corresponding to rhesus macaques of Chinese and Indian origin. Supplementary Fig. S4 shows an additional PCA analysis comprising only *M. mulatta* individuals, again showing a clear distinction between the two main macaque populations^6,20–23^ with putative hybrid individuals^24^ clustering in between. Although *M. sylvanus* should serve as an outgroup to the other three species, the PCA includes only a single sample, which is insufficient to form a distinct cluster.

Lastly, to investigate the relationship between alignment specificity to the human reference and within-species nucleotide diversity across primates, we classified regions of each of the 47 primate reference genomes into three categories: regions aligning uniquely to the human genome, regions aligning to multiple human loci, and regions with no detectable human homology. For the autosomes, the average diversity was generally lowest in regions that align uniquely to the human reference (1.6×10^-3^), while higher diversity was observed for regions that have non-unique (1.85×10^-3^) or no homology (1.95×10^-3^) to the human reference (Fig. 5A). The mean enrichment of diversity within regions with no homology was 1.27× (minimum of 0.82× and maximum of 2.46×) compared to uniquely aligning regions, and 1.07× (minimum of 0.39× and maximum of 1.76×) compared to non-uniquely aligning regions. This is in line with the expectation that uniquely aligned regions tend to be conserved and thus have slower evolutionary rates compared to more species-specific regions. Interestingly, while X chromosome diversity was lowest in uniquely aligning regions (8.32×10□□), regions aligning to multiple human loci had slightly higher diversity (1.15×10□³) compared to regions with no homology to the human reference (1.13×10□³), especially in primate families more distantly related to humans (Fig. 5B). This may be a consequence of the unique evolutionary dynamics of duplicated regions on the X chromosome^25,26^, but should be interpreted with caution given that the amount of X chromosome sequence available for alignment was lowest in non-unique regions (ranging between 0.93 and 9.52 Mb of total alignable sequence).

**Figure 5.**
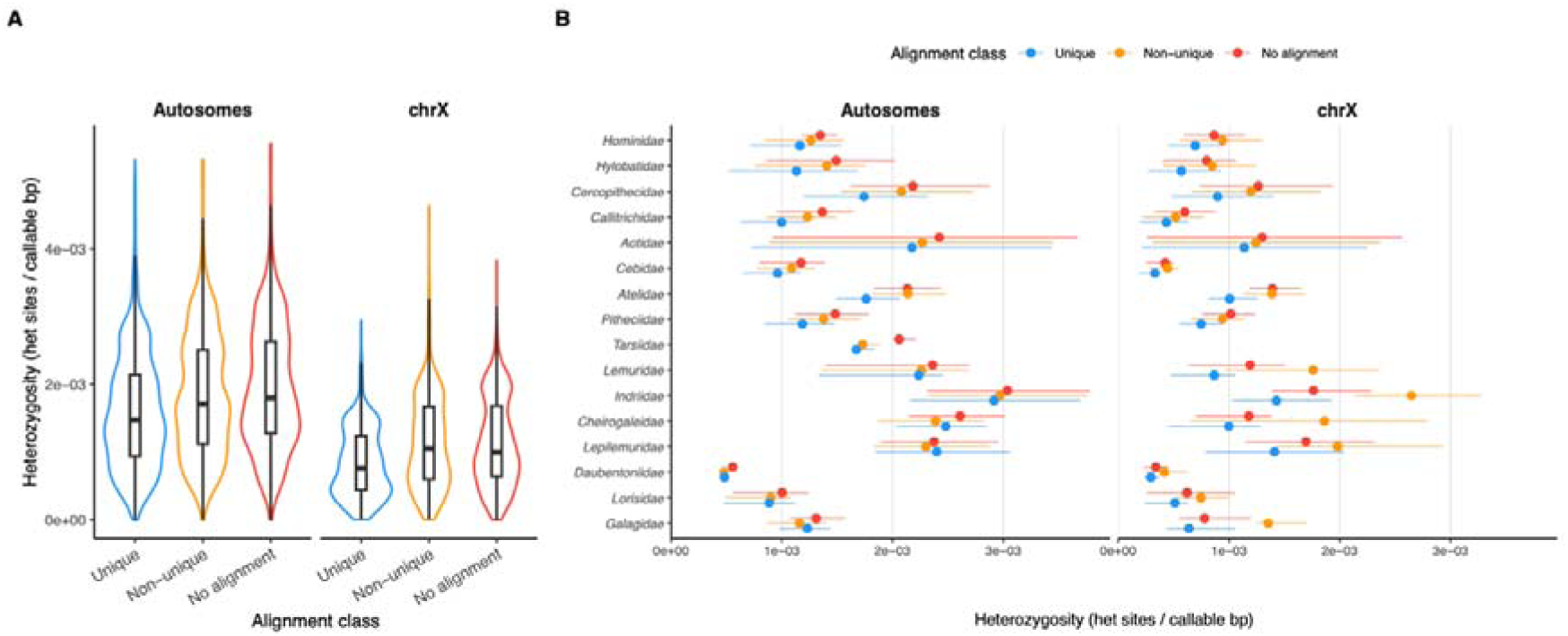
Patterns of genomic diversity across three different classes of sites with respect to primate reference alignability to the human sequence. **A.** Distribution of heterozygosity across all individuals for the three region classes and two chromosome types. **B.** Estimates of mean heterozygosity (points) with the corresponding interquartile range (whiskers) across individuals within each of the 16 primate families, for the three region classes and two chromosome types.

## Conclusions

This study establishes a comprehensive genome-scale nucleotide diversity panel of non-human primates, encompassing the largest number of individuals to date. By incorporating ploidy-aware variant calling and high-quality genome assemblies, the resource overcomes previous limitations in primate comparative genomics. The resulting dataset and multiple-genome alignment provide a robust foundation for advancing research in primate evolution and conservation.

## Methods

### Raw Data Acquisition and Preparation for Read Mapping

Paired-end short-read sequencing data were downloaded from the FTP server of the Sequence Read Archive (SRA; ftp://ftp.sra.ebi.ac.uk/vol1/) as already separated FASTQ files. We queried the NCBI Sequence Read Archive (SRA) database using the Biopython Entrez module^27^ to identify whole genome sequencing datasets for each target genus. Search queries were constructed using genus names as organism identifiers and filtered to include only datasets generated through whole genome sequencing (WGS) or whole genome amplification (WGA) strategies. Retrieved metadata summaries were manually curated to retain samples consistent with our study objectives, primarily those derived from wild-caught individuals or population diversity panels, where possible. The code for querying NCBI is available on the project-associated Github repository (https://github.com/Bjarke-M/Primate-Recalling). In total, we processed data from 3,240 individual biosamples, spanning 269 species and 71 genera. A complete list of BioSample IDs and short-read run accessions with corresponding download links is available in Supplementary Table S4.

If an individual BioSample contained multiple run accessions of short-read data, the corresponding FASTQ files were concatenated using the cat program (v8.32)^28^. All FASTQ files were processed into unmapped BAM (uBAM) files using the FastqToSam program from the picard suite of programs (v3.1.1)^29^ prior to read mapping.

A total of 47 reference genome assemblies used for mapping and the human T2T reference were obtained from the FTP server of the National Centre for Biotechnology Information (NCBI; https://ftp.ncbi.nlm.nih.gov/) and the DNA Zoo^30^ repository (https://dnazoo.s3.wasabisys.com/index.html?prefix<u>=</u>). Additionally, the genome assembly of *Tupaia belangeri* used as an outgroup in the multiple-genome alignment was obtained from http://tupaiabase.org. A complete list of reference assemblies with corresponding download links is available in Supplementary Table S2.

Short reads and assemblies were downloaded using wget (v1.21.1)^28^. Prior to short-read mapping, each reference assembly was modified to contain only contigs ≥1000 base pairs (bp), using the reformat.sh script of the BBmap suite of programs (v38.18)^31^. Additionally, masking of Y-linked pseudoautosomal regions in great ape (*Pan troglodytes*, *Pan paniscus*, *Gorilla gorilla*, *Pongo pygmaeus* and *Pongo abelii*) and the siamang (*Symphalangus syndactylus*) genome assemblies was done using the bedtools maskfasta program (v2.31.1)^32^. The assemblies were indexed using bwa index (v0.7.17)^33^ and samtools faidx programs (v1.19)^34^, and the corresponding sequence dictionary was created using the Genome Analysis Toolkit (GATK v4.4.0.0)^5^ CreateSequenceDictionary program.

### Read Mapping and Preparation for Variant Calling

To parallelize read mapping, each uBam file was further split into shard files containing 48,000,000 read pairs using the picard SplitSamByNumberOfReads program. Adapter sequences were marked in each shard file using the picard MarkIlluminaAdapters program, followed by extraction of read sequences using the picard SamToFastq program, which were piped into the bwa mem program used for read mapping. The output of bwa mem was finally piped into the picard MergeBamAlignment program, producing a mapped shard BAM file. Finally, the shard files were concatenated into individual-specific mapped BAM files using the picard MergeSamFiles program. We marked duplicated reads in each individual-specific BAM file using the picard MarkDuplicates program, followed by sorting of the BAM file by the corresponding reference sequence coordinates using the picard SortSam program. Individual-specific read coverage of each reference contig was calculated using the samtools depth program. The total read count and the count of mapped reads were estimated using samtools flagstats, and the samtools coverage was used to estimate per-contig average mapping quality. Genome-average mapping quality was calculated as the mean of the per-contig values weighted by the contig length.

Calling sex chromosome-derived contigs with correct ploidy required identifying such contigs in the reference genome and knowing the sex of the sequenced individuals. Both types of information were not available for some of the reference genomes/samples. We therefore took several steps to fill in these gaps. First, for each reference genome except *Cephalopachus bancanus* which is highly fragmented (N50 = 5.3 Mb), we identified a single contig with the largest number of nucleotides aligned to the human X chromosome according to the Multiple Alignment Format (MAF) file generated from the multi-genome alignment. Next, we used the coverage of these X chromosome-derived contigs (one per reference genome) normalized by the genome-wide coverage of the same sample to perform genetic sexing of samples with genome-wide coverage above 10× and fraction of the whole genome with non-zero coverage above 75%. We labeled samples having such normalized coverage above 0.8 as females and below 0.7 as males forming a set of confidently sexed individuals. Samples with intermediate values of normalized coverage underwent additional analyses before assigned a sex.

We next used the set of confidently sexed individuals to search for additional contigs within the respective reference sequence with X and Y chromosome characteristics. To do so, we first estimated the normalized coverage (relative to the genome-wide value) and the fraction of positions with non-zero coverage for each contig in each sample and then took the median for each contig across all samples mapped to a particular reference, separately for males and females. Next, we applied the following criteria in this order: a) if a contig was annotated as Y chromosome, we kept this annotation; b) if the ratio of median fractions of the contig’s length with non-zero coverage between males and females was larger than 1.5, we annotated the contig as Y chromosome; c) if at least 5% of the contig’s length was aligned to the human X chromosome and the median relative coverage in males was below or equal to 0.75 while being above 0.75 in females, we annotated the contig as X chromosome. For all contigs selected based on criteria (b) we additionally checked if the fraction of the contig’s length covered was higher in males than in females across different species, as this measure can be affected by sex but also by the phylogenetic distance to the reference. Results were consistent with annotating these contigs as Y chromosomal in all cases except for the *Lemur catta* reference. In the latter case, only contig NC_059156.1 was annotated as chromosome Y. For two references, namely *Callimico goeldii* and *Piliocolobus tephrosceles*, there were only females and only males mapped against them, respectively. In these two cases, we only tested the median coverage for the sex that is available to annotate X chromosome contigs using criterion (c). All identified contigs are available in Supplementary Table S3.

After identifying the X chromosomal and, where available, Y chromosomal contigs in each reference, we characterised each sample with the coverage and fraction of the length with non-zero coverage separately for the X chromosome (combining all the X chromosomal contigs), Y chromosome (combining all the Y chromosomal contigs) and autosomes (combining all the remaining contigs). We used these characteristics to identify sex for the low coverage samples (genome-wide coverage below 10x or/and less than 75% of the genome with non-zero coverage). For samples mapped against a reference with no Y chromosome, we assigned female sex if the ratio of X chromosome coverage to autosomal coverage was above 0.8, male sex if this ratio was below 0.7 and kept the sex unassigned for intermediate values. We note, however, that for samples with very low coverage, this ratio converges to 1 for both sexes, resulting in treating all such samples as females in subsequent analyses. Hence, for samples mapped against a reference with a Y chromosome, we instead used a ratio of the fraction of the Y chromosome length with non-zero coverage over the same fraction for the X chromosome, which is more informative about sex for low-coverage samples. As the fraction of a Y chromosome-derived contigs’ length that can be covered with reads from the female genome will differ depending on which regions of the Y chromosome were represented by the contig we treat as Y chromosome, there is no universal threshold that can be used for all references. Instead, we used a nearest neighbour classifier with three neighbours trained on the high-coverage samples. Finally, for both high and low coverage samples with the relative X chromosome coverage between 0.7 and 0.8 we used the sex reported in metadata if available and if not labeled samples with relative coverage above 0.75 as females and as males otherwise.

### Variant Calling and Genotyping

We parallelized variant calling by extracting 2 megabase (Mb) genomic intervals from individual-specific mapped BAM files using the samtools view program, followed by assigning a new read-group to each region-specific BAM using the GATK AddOrReplaceReadGroups program. Each region-specific BAM was used as input to the GATK’s HaplotypeCaller program, with the -ploidy option set to reflect the genetic sex of the individual and the context of the region (autosomal, X-linked, Y-linked or mitochondrial). HaplotypeCaller was run using the option -ERC BP_RESOLUTION to generate gVCF output files that contain a call for every site of the reference genome (*i.e.* both variant and non-variant sites were emitted) for each 2 Mb region and each individual. The output gVCFs were indexed using GATK’s IndexFeatureFile program.

GATK’s GenomicsDBImport program was used to merge region-specific, per-individual gVCFs across individuals of the same species into a species-specific GenomicsDB database used for joint genotyping, conducted using GATK’s GenotypeGVCFs program. The resulting region-specific, multi-individual genotyped gVCFs were concatenated to produce chromosome-specific gVCFs, or multiple-contig gVCFs for contigs less than 30 Mb in length, using the picard SortVcf program. Concatenated gVCFs were indexed using GATK’s IndexFeatureFile program.

### Callability Mask and gVCF Filtering

We used bcftools^35^ v1.22 to manipulate gVCF files. To filter the gVCFs to keep only reliable genotypes, we first created a species-specific callability mask. For this, we used samples with sequencing depth of at least 10× that passed a species-specific cutoff for the fraction of the genome covered with at least one read for both autosomes and chromosome X (Supplementary Table S1). Then, for each species, we summarised coverage for each position in the genome across this subset of samples and calculated the average across all autosomal and chromosome X-derived contigs separately. Next, for each contig, we kept positions with coverage within 0.5× and 2× of the average coverage of the corresponding contig type (autosomal or chromosome X derived) and merged these positions into bed regions. Finally, we created a positive callability mask by keeping regions at least 100 bp long.

Next, we applied these masks to the gVCF files. We kept only SNPs and converted multiallelic variants into separate biallelic records. Additionally, we excluded positions with the ‘LowQual’ flag, or meeting at least one of the following criteria: QD (quality by depth) < 2; FS > 60; MQ < 40; SOQ > 3; ReadPosRankSum < -8.0; MQRankSum < -12.5. Individual genotypes were set to missing if meeting at least one of the following criteria: DP (depth) < 5; GQ (genotype quality) < 30; AD (allelic depth) < 3 for heterozygous positions; ratio of depth for the allele with lower depth to the depth for the allele with higher depth < 0.3. The resulting files were used when estimating heterozygosity and other genomic statistics. Only contigs at least 1 Mb long were used.

### Sample Quality Control

For quality control, we used the following sample characteristics:

A. Fraction of the reference genome covered with at least one read (*fraction covered*);
B. Sequencing depth for the covered fraction of the reference genome (*sequencing depth*);
C. Missing rate, defined as the count of ./. genotypes divided by the length of the species-specific callability mask (*missing rate*);
D. Divergence from the reference sequence, defined as the count of 1/1 genotypes plus half the count of 0/1 genotypes divided by the length of the species-specific callability mask (*sequence divergence*).

For *fraction covered*, we set a lower threshold equal to 0.9 for both autosomes and chromosome X for most genera. Thresholds lower than 0.9 and or different thresholds for autosomes and chromosome X were set for *Arctocebus*, *Lepilemur*, *Loris*, *Microcebus*, *Mirza*, and *Perodicticus* after a visual inspection of the relationship between *fraction covered* and *sequencing depth*.

If a sample had *fraction covered* or *sequence divergence* too high or too low compared to other samples within the same species cluster with respect to the reference species, it might indicate compromised sequencing quality or a mismatch between the observed phylogenetic distance to the reference and the species label. To identify such outliers, we first grouped species into clusters of species phylogenetically equidistant to the reference species. Species with clear population structure (*Cercopithecus mitis, Macaca mulatta, Papio anubis, Pan paniscus* and *Pan troglodytes*) were further split into subclusters based on species-specific thresholds for *sequence divergence*. Next, for each (sub)cluster, we created a subset of high-quality samples. These are samples with a) *fraction covered* above the threshold for both autosomes and chromosome X; b) *sequencing depth* above 20; and c) *missing rate* below 1%. For each such subset, we defined the median and interquartile ranges (IQR) for *fraction covered* and *sequence divergence* based on autosomal contigs. We then described every sample, including those that did not meet the above-mentioned criteria for high quality, with the difference between the sample’s value and the median of the sample’s (sub)cluster, divided by the interquartile range of the (sub)cluster. We then use these two IQR-based robust z-scores to identify outliers that have a) too low *fraction covered*, b) too high or c) too low *sequence divergence* based on thresholds described in Supplementary Table S1. We then assigned samples an ‘ok’, ‘warn’ or ‘fail’ status based on these characteristics (see Supplementary Table S1 for details). Finally, this status was manually modified for 67 samples based on visual inspection and / or additional information as described in Supplementary Table S1.

### PCA data filtering

For the PCA analysis, all samples with less than 20× coverage were not considered. Only biallelic sites and contigs at least 1Mb in length were used. Sites with a minor allele frequency less than 10% were filtered out. Variants with linkage disequilibrium of more than 0.5 (Lewontin’s D)^36^ in 1000bp windows were filtered out. Contigs identified as X-linked were used for the chromosome X PCA, while autosomal contigs were used for the corresponding autosomal PCA analysis. Although the underlying dataset encodes the X chromosome as haploid in males, we treated male X-linked genotypes as diploid for PCA, since the analysis required all samples to have the same ploidy. All filtering steps were done with bcftools, as bcftools view -Ou -s {samples} | bcftools view -Ou -m2 -M2 | bcftools view -Ou -i ‘MAF[0]>=0.1’ | bcftools +prune -Ob -m LD=0.5 -w 1000bp --random-seed 10.

### Multiple-Genome Alignment

To facilitate between-species comparisons, we generated a multiple-genome alignment comprising 48 primate reference assemblies (47 reference genome assemblies used for mapping and the human T2T reference) and the outgroup *T. belangeri*. Specifically, for each reference genome, we kept only contigs longer than or equal to 1 Mb in length and applied RepeatMasker^37^ v4.1.8 with Dfam^38^ v3.9 repeat library to those sequences that were not already soft-masked. Next, we ran progressive cactus^9^ v2.9.9 using a published Snakemake workflow^39^. The resulting alignment in the HAL format was converted to MAF format with the ‘cactus-hal2maf’ function with the human genome as the reference. This maf-formatted alignment was then processed with taffy provided within the cactus container to calculate the fraction of the human genome covered by each of the remaining 48 genomes, as well as pairwise divergence from the human sequence.

### Workflow Management and Data Availability

All steps of the bioinformatic pipeline were implemented within the gwf workflow management framework (v2.0.5)^40^ can be found at https://github.com/Bjarke-M/Primate-Recalling.

The individual-specific mapped BAM files (in CRAM format), multi-individual genotyped gVCFs at base-pair resolution as well as filtered versions of the data (variant only VCFs and filtered BCF files) have been uploaded to the Electronic Research Data Archive (ERDA) at Aarhus University (AU, Denmark) and can be found at https://pure.au.dk/portal/en/datasets/primate-diversity-panel/.

## Acknowledgements

All of the computing for this project was performed on the GenomeDK cluster (https://genome.au.dk/). We would like to thank GenomeDK and Aarhus University for providing computational resources and support that contributed to these research results. We also thank Jesper Lykkegaard Karlsen for help with data upload and management in ERDA.

## Supplementary Figures

**Supplementary Figure S1.**
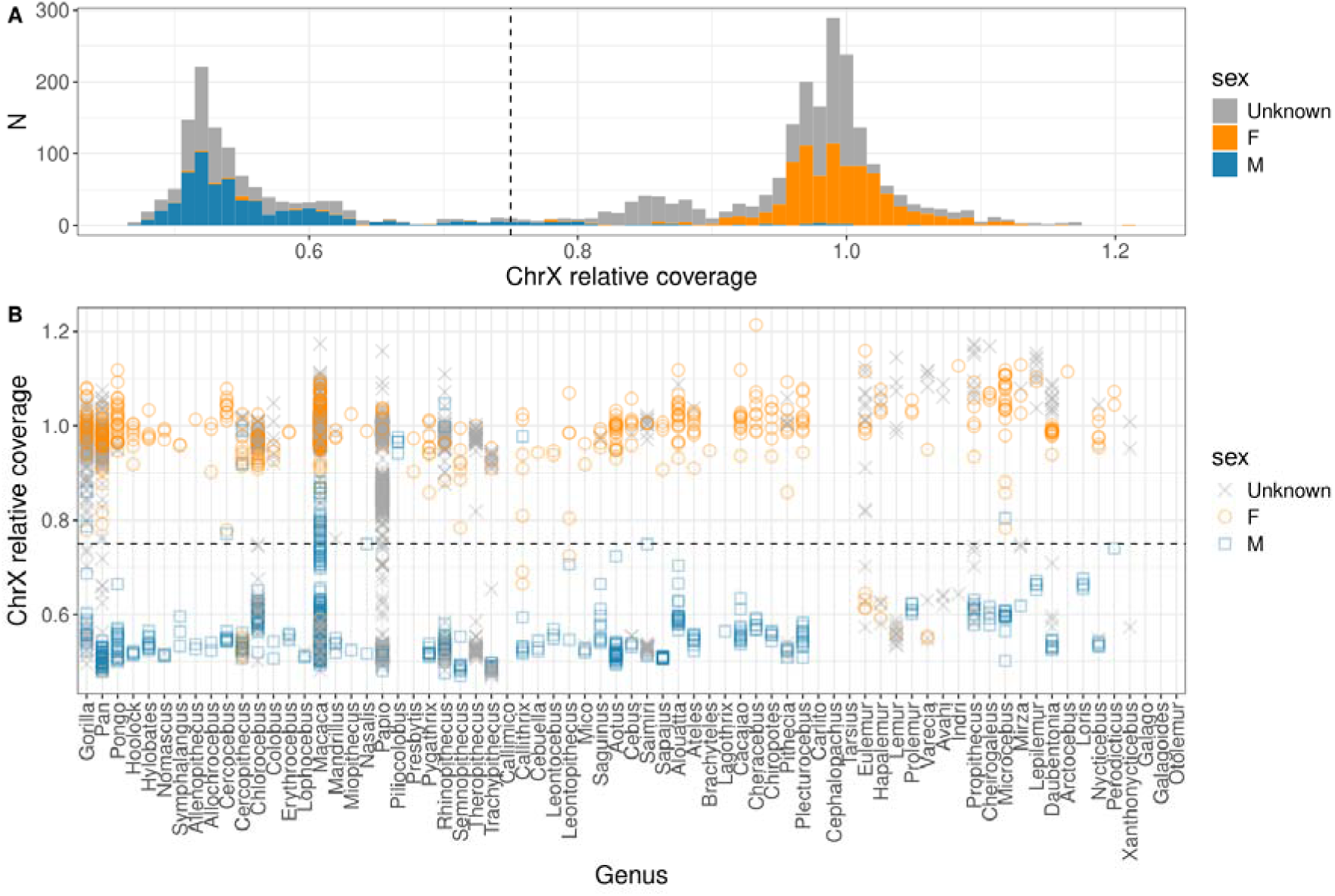
Genetic sexing of samples. Relative chromosome X coverage (sequencing depth of chromosome X divided by the sequencing depth of the autosomes). For both chromosome X and autosomes, only contigs longer than or equal to 1 Mb are considered. Panel **A.** displays the distribution in the entire dataset, while panel **B.** shows individual samples grouped by genus. Color shows sex as reported in the metadata. The dashed line marks the 0.75 cutoff used for genetic sexing. In both panels, sample ‘SAMEA80831668’ is excluded as it was sequenced with chromosome Y capture and as a result has a relative chromosome X coverage of 6.843 due to homology between the sex chromosomes.

**Supplementary Figure S2.**
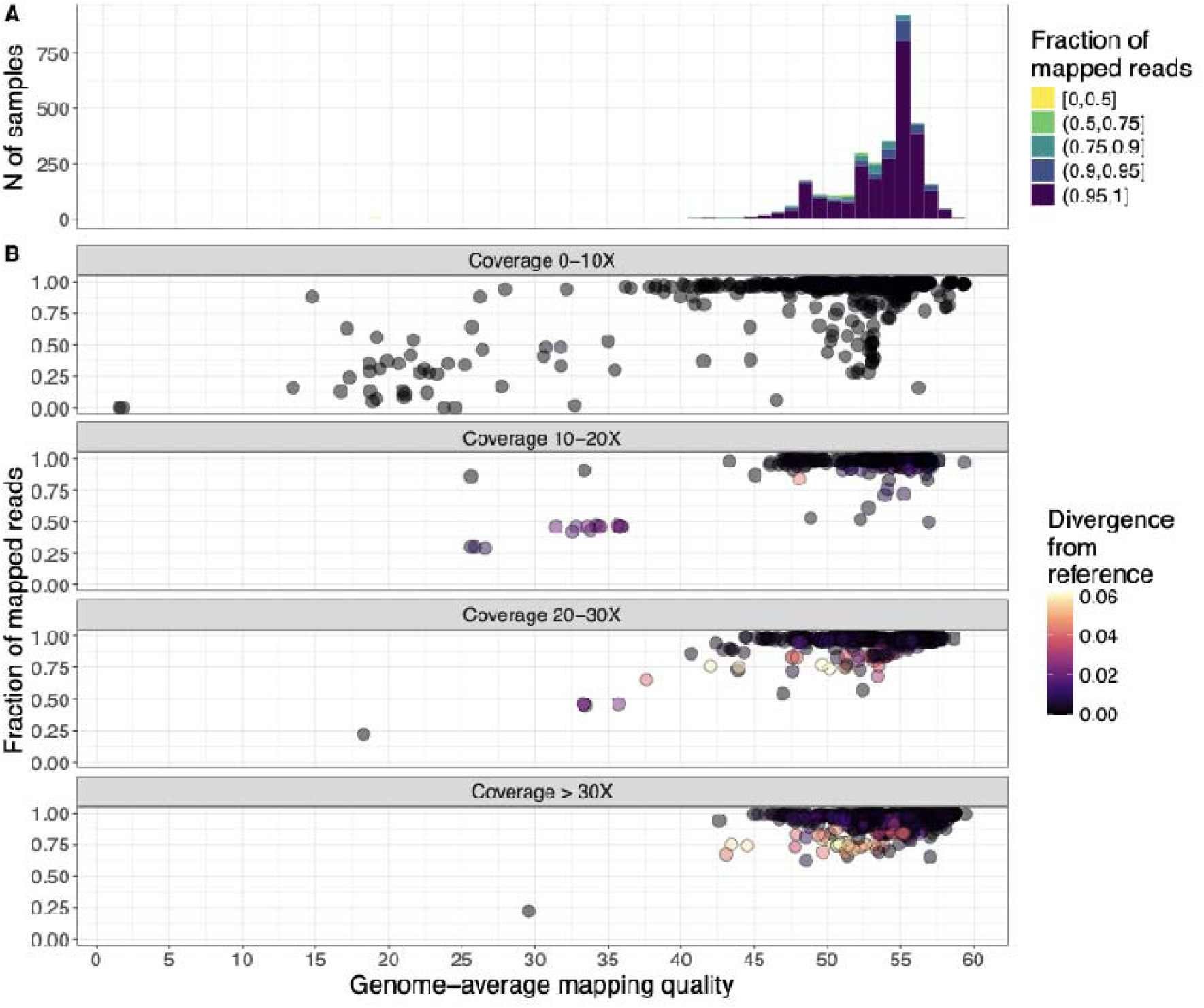
Sample distribution based on average read-mapping quality and the fraction of the reads mapped to the reference genome. A. Number of samples with different mapping quality and fraction of mapped reads. B. Relationship between coverage and divergence from the reference versus mapping statistics.

**Supplementary Figure S3.**
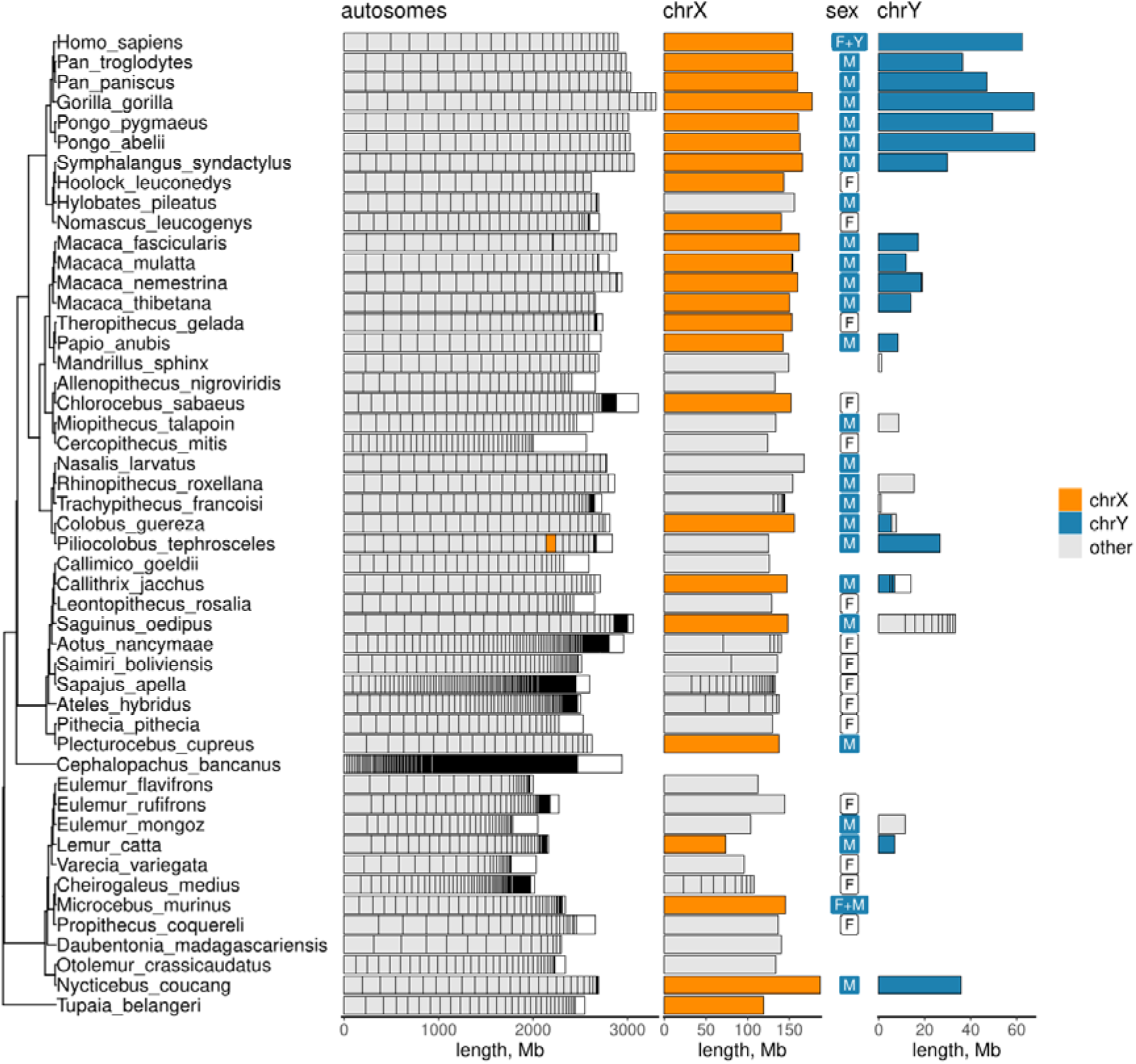
Reference genomes’ length. For each of the 47 reference genomes (plus the human genome and the genome of the outgroup *T. belangeri*), we show the total length of the autosomal as well as chromosome X- and chromosome Y-derived contigs. Each rectangle represents a single contig. Contigs shorter than 1 Mb are combined, and their total length is represented by a white rectangle on the right part of the bar. The colour of a rectangle represents its annotation in the reference genome. Sex chromosome-derived contigs were identified as described in the Methods. This analysis was only done for contigs longer than 1 Mb. Note that a contig annotated as chromosome X in the genome of *P.tephrosceles* is re-classified as autosomal based on our analysis, while a different contig is identified as being chromosome X-derived. Sex of the sample used to produce the reference is reported where available. F - female, M - male, F+M - a pooled sample of a male and a female was used, F+Y - a female sample was used, and chromosome Y sequence from a different individual was added to the assembly.

**Supplementary Figure S4.**
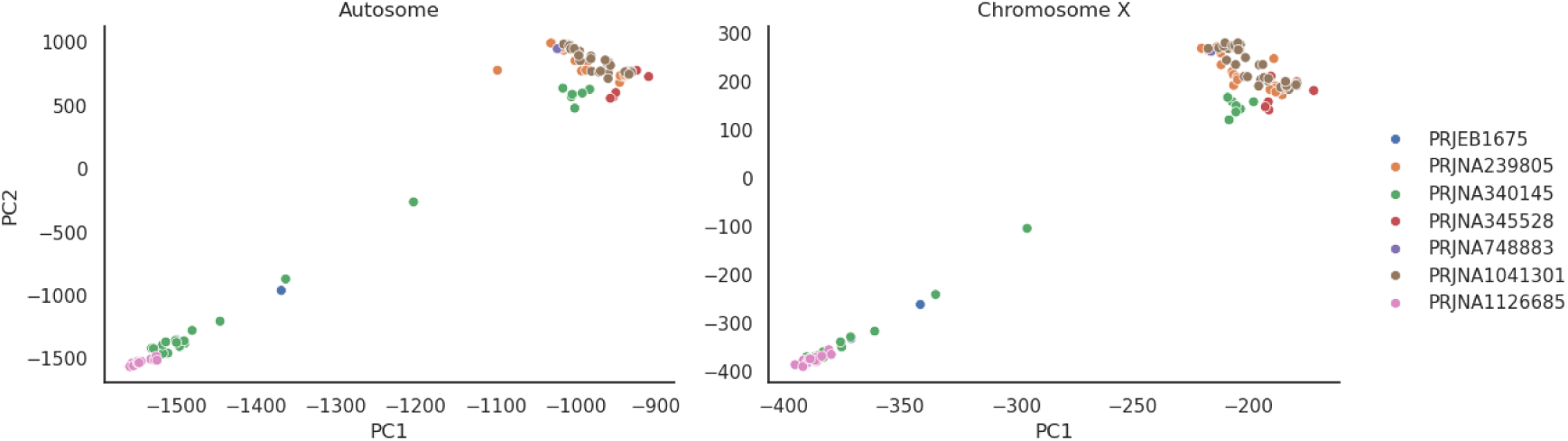
PCA of *Macaca mulatta* individuals mapped to the *Macaca mulatta* reference assembly based on the autosomal and the X-linked variants. The top-right cluster corresponds to the Chinese *Macaca mulatta* population, with most of the individuals sourced from BioProject PRJNA1041301. The lower cluster corresponds to BioProject PRJNA1126685, which is from a free-ranging, Indian-origin population of Cayo Santiago *Macaca mulatta* in Puerto Rico. The intermediate macaques are mostly from BioProject PRJNA340145 and colored in green, comprising individuals from both the Indian and Chinese populations, as well as two hybrid individuals.

